# A genetic perspective on the recent demographic history of Ireland and Britain

**DOI:** 10.1101/2024.03.08.584042

**Authors:** Ashwini Shanmugam, Michael Merrigan, Seamus O’Reilly, Anne M. Molloy, Lawrence Brody, Orla Hardiman, Russell McLaughlin, Gianpiero L. Cavalleri, Ross Byrne, Edmund Gilbert

**Author notes:** Equal contribution (Ross Byrne, Edmund Gilbert).

## Abstract

**Background:** While subtle yet discrete clusters of genetic identity across Ireland and Britain have been identified, their demographic history is unclear.

**Methods:** Using genotype data from 6,574 individuals with associated regional Irish or British ancestry, we identified Irish-like and British-like genetic communities using network community detection. We segregated Identity-by-Descent (IBD) and Runs-of-Homozygosity (ROH) segments by length and approximated their corresponding time periods. Through this, we inferred the regional Irish and British demographic histories in these time periods by (1) estimating genetic relatedness between communities, (2) estimating changes in effective population sizes, (3) inferring recent migration rates across Ireland and Britain, and (4) estimating changing affinities to regional European populations. For a subset of the Irish communities, we determined the enrichment/depletion of surnames within the genetic communities.

**Results:** Through patterns of IBD-sharing and ROH, we find evidence of recent population bottlenecks in the Orcadian, Manx and Welsh communities. While the structure in Ireland is subtler, the communities share relatively more IBD segments that are shorter in length, and the genetic differences between the Irish communities are more subtle on average, when compared to the British communities. Regional effective population size trajectories indicate a similar demographic history throughout the island of Ireland. Further, we observe a stable migration corridor between north-east Ireland and south-west Scotland while there is a recent migration barrier between South-Eastern Ireland and Western Ireland. We observed an enrichment of Anglo-Norman and English surnames in the Wexford community while within the West Ulster-Argyll community, we saw an enrichment of Gallowglass and Scottish surnames.

**Conclusions:** Using well-annotated Irish and British reference genotypes, we observed temporal changes in genetic affinities within and between genetic communities in Ireland and Britain. In addition, using effective population size estimates and levels of haplotype-sharing, we detected varying degrees of genetic isolation in some Irish and British genetic communities across time. Using these new insights into the regional demographic history of Ireland and Britain across different time periods, we hope to understand the driving forces of rare allele frequencies and disease risk association within these populations.

## INTRODUCTION

The subtle but distinct fine-scale population structure of Ireland is well-characterised^1–3^. The broad population structure in Ireland segregates along historical provincial boundaries^2,3^. There is a general east-to-west and a north-to-south gradient of British ancestry with higher proportions observed in northeast and southeast Ireland^3^, consistent with demographic and political history of the country^4,5^. Furthermore, studies have investigated the ancestry profiles of Ireland, with a focus on Norwegian ancestry. Studies using Y-chromosome markers^6,7^ found little evidence of an enrichment of Norse/Scandinavian haplogroups in Irish males with putatively Norse surnames while others, using genotypes from modern European populations to capture ancestry proportions population^2,3,8^, found conflicting evidence of Norwegian ancestry in the Irish population. Recently, ancient genomes from various timepoints have been used to illustrate that while there was indeed Norwegian Viking ancestry in Ireland, there was also significant gene flow from Ireland and the British Isles^9^, into Scandinavia during the Viking age^10^. However, the genetic footprints of the demographic history within the island of Ireland is relatively under-studied — there is a gap in our understanding of how the effective population size and genetic structure within regions of Ireland has changed over time.

The demographic history of a population is reflected within the genetic variation of its current generation^11–13^. Past population sizes affect haplotype diversity and patterns of variation which in turn may contribute to relative risk of complex traits^14,15^. By understanding population structure and the demographic events that shaped it, we can effectively control for their effects while investigating the dynamics of selection and the association of genotypes to phenotypes within a population^16,17^. Insights into recent demographic history can also provide vital information about the distribution of rare functional variation as long-range haplotypes tend to be more recent and can therefore be used to impute rare variant burden in samples to reveal novel genotype-phenotype associations in complex traits and diseases^18–20^. To understand the patterns of genetic variation and the forces that drive it in a population such as Ireland, it is vital to study the effects of demographic events such as migrations^9,21,22^, to estimate the extent of admixture, and population contractions - e.g. due to famine^22^.

A variety of approaches are available to investigate demographic histories, typically based on one of three broad approaches; (1) modelling patterns of genetic variation using allele frequency spectra or density of heterozygous sites in a region^23^, (2) measuring decay of linkage equilibrium at sites which are identical-by-state (IBS)^24,25^, or (3) leveraging patterns of long-range haplotype-sharing or identity-by-descent (IBD) segment sharing^26–29^. However, due to their properties, IBD segments are best suited to applying principles of coalescence to infer features of a population’s recent demographic history^14,15^ (typically within the past approximately 200 generations^28^). Since successive recombination events break haplotypes during meiotic transmissions, the length of an IBD segment is a function of time to a common ancestor while the number of segments are direct traces of recent coalescence events^30^. Therefore, IBD segments have been leveraged to provide comprehensive demographic histories of populations by (a) detecting the extent of genetic relatedness (i.e. population structure) defined by an IBD-sharing network^31–33^, (b) inferring trajectories of effective population size over time^28^, (c) identifying migration corridors and barriers^34^.

In this context, our study aimed to: (1) detect the extent of background relatedness in British and Irish regions, using long haplotype sharing, (2) identify the different regional demographic histories in Ireland and Britain, (3) leverage long haplotype sharing to disentangle European ancestry in Ireland and Britain.

## RESULTS

### Population structure in Ireland and Britain

We inferred fine-scale population structure from the largest collection of Irish reference haplotypes (*n* = 3,502) with geographic provenance assembled to date combining datasets of Irish and British ancestries (Table 1). Identifying genetic communities based on haplotype sharing patterns allowed us to study regional demographic histories of Ireland and Britain. We identified a total of 25 genetic communities using the Leiden network community detection algorithm^35^ over three levels of recursive clustering, constructing a network of summed IBD-segment sharing amounts between individuals (see Methods).

**Table 1:** Summary of datasets.

| Dataset | Population | Geographic Data |  |
| --- | --- | --- | --- |
|  |  | Present | Absent |
| ALS Case Control Cohort | Northern Ireland | 7 | 0 |
|  | Republic Of Ireland | 533 | 445 |
| Irish DNA ATLAS | Northern Ireland | 24 | 0 |
|  | Republic Of Ireland | 166 | 5 |
| People of the British Isles | England | 2405 | 29 |
|  | Isle of Man | 56 | 0 |
|  | Northern Ireland | 69 | 0 |
|  | Orkney | 136 | 1 |
|  | Republic Of Ireland | 6 | 1 |
|  | Scotland | 152 | 30 |
|  | Wales | 290 | 5 |
| TRINITY Student Study | Republic of Ireland | 0 | 2214 |

Confirming previous reports^2,3^, these genetic communities segregated by geography and were assigned labels to broadly reflect these geographic affinities (Supplemental Table 1). The first recursion classified individuals into two communities (Supplemental Fig. 1a), one with predominantly Irish membership and the other predominantly with British membership. The second recursion largely split the communities by historical or administrative boundaries — provinces within Ireland (n = 3) and generally countries within the UK (n = 5) (Fig. 1a-b). The third recursion identified fine-scale structure within these groups (Supplemental Fig. 1b-i). We identified previously unobserved genetic structure within Ireland (N.Kerry, S.Leinster, three Dublin communities) and the UK (East Anglia).

**Fig. 1.**
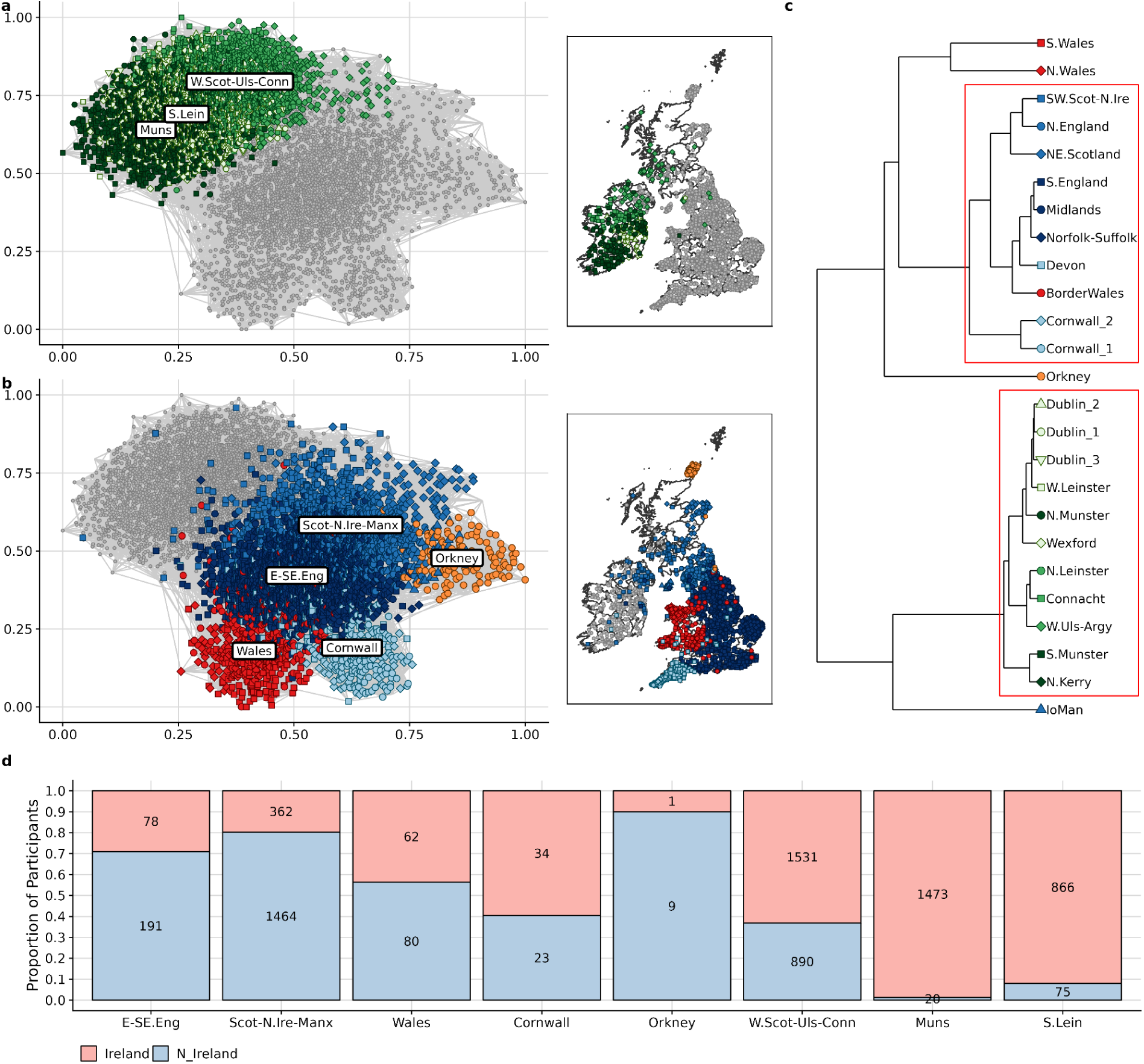
Genetic Structure in Ireland and Britain. (a & b) A graphical projection of the network of genetic communities detected in Ireland and Britain. Every point on the plot is an individual with British or Irish ancestry and the colours of the points indicate genetic community membership. Each line, or edge, is the average length of IBD segments shared between a pair of individuals with line thickness corresponding to the total length of segments shared. For each individual, we show the top 10 edges for plotting. The maps adjacent to the networks show the geographic origin of individuals (see Methods for more details). (c) hclust dendrogram of 3rd level genetic communities showing genetic similarity. The red rectangles around the branches indicate high certainty grouping and the colour of the points indicates the 2nd level genetic community and shape of the points indicate the 3rd level genetic community. (d) Barplot depicting the predicted regional ancestry of Irish and N.Irish participants from the UK Biobank. The x-axis shows the second-level genetic communities within Ireland and Britain to which samples were assigned by our classifier. The colours in the stacked barplot indicate the reported birthplace of the UK Biobank participants and the y-axis displays the proportion of the samples assigned to the regional ancestry group from each birthplace.

We further characterised this fine-scale structure by performing hierarchical clustering using *pvclust*^36^ on the average inter-community haplotypic lengths^37^ (see Methods). We find supporting evidence that these communities represent meaningful divisions in our sample of Irish and British haplotypes (Fig. 1c and see Supplemental Note 1 for full analysis). Permutations of total-variation distances (TVD)^21^ (see methods) confirmed the robustness of the communities (p-value << 0.01, Supplemental Table 2).

We sought to demonstrate the value of these findings in the context of the UK Biobank (UKB), one of the largest accessible datasets of Irish and British ancestry^38^. We demonstrate that the UKB self-reported “white British” or “white Irish” ethnicity label poorly captures the diversity of Irish ancestry (Fig. 1d). We first randomly split our Irish and British dataset 7:3 as training and validation datasets. Using the training set and their cluster labels, we trained a Naive Bayes classifier to predict regional Irish and British ancestry. The median accuracy of this model was 66.9% and was assessed using the validation dataset. We applied the model to predict the regional ancestry of the UKB participants reporting a Irish and Northern Irish birthplace (“UKB Irish”) (n = 7,159; see Methods and Supplemental Table 3, Supplemental Fig. 2a-d). We find that 67.81% of these Irish or Northern Irish-labelled UKB participants are predicted to have regional ancestry most commonly found in the Republic of Ireland. Breaking down by broad region defined by our second-level clusters (Fig. 1), 33.82% of UKB Irish are classified as of Western/North-Western Irish ancestry, 20.85% of Southern Irish ancestry and 13.14% of Eastern Irish ancestry. Of the remaining 32.19%, 6.68% were classified into one of any other British-majority ancestry cluster, and 25.51% were classified in the Northern Irish/South-West Scottish/Manx-like group (Supplemental Table 3). This group was predominantly made up of those UKB Irish born in Northern Ireland (80.17% vs 19.83% who were born in the Republic of Ireland). Our findings show that the UKB self-reported “white British” or “white Irish” group is biassed in terms of a sampling scheme, being weighted towards the north of the island which accounts for nearly 60% of the predicted Irish-like participants. While the UKB is a large repository of Irish ancestry, it is only with the use of additional well-annotated reference datasets that any fine-scale understanding of regional ancestry within Irish samples from this dataset is possible.

#### Genealogy and population structure

Genealogical data can also provide additional context to genetic communities detected through haplotype analysis. Genealogical data available through the Irish DNA Atlas allowed us to test the preponderance of different surname origins (i.e., English, Scottish, Irish) within each of the Irish third-level clusters (Fig. 2). We observed a significant enrichment of Scottish and Gallowglass surnames in the SW.Scotland-N.Ireland community — matching historical settlement of Scottish mercenaries in that area between the 13th and 16th centuries^4^. Within the N.Munster and N.Kerry communities, we observe an enrichment of Welsh surnames. In Wexford, we find an increase of English, Anglo-Norman and Scandinavian surnames which possibly may reflect older Anglo-Norman settlements in these regions^5^. When we extended this analysis to individual surnames (Supplemental Fig. 3), we observed an enrichment of “Walshes” in N.Kerry and W.Leinster, “Sullivans” and their variants in N.Kerry and S.Munster, “Ryans” in N.Munster, “O’Donnells” in W.Ulster-Argyll.

**Fig. 2.**
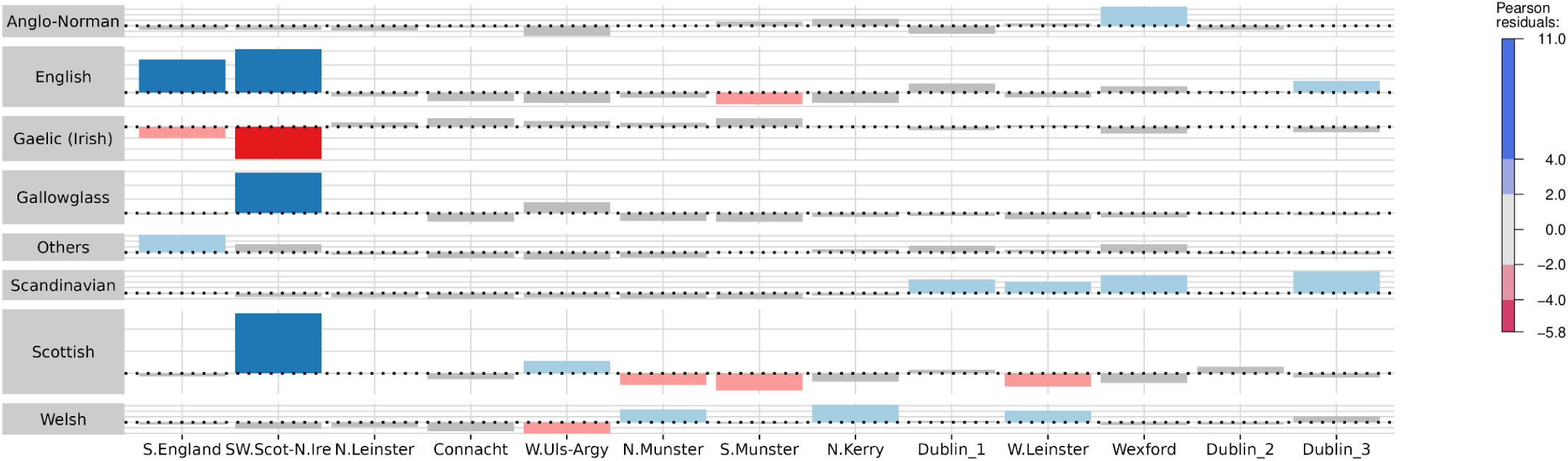
Surname association with regions in Ireland. Using a Chi-square test, we estimated the differential occurrence of surnames (classified according to their origins into 8 categories) within the Irish genetic communities. The colours of the bars indicate the Pearson residual value of the surname origin class (along y-axis) within each genetic community (x-axis).

### Demographic Profiling of Genetic Communities Across Ireland and Britain

With a set of regional genetic communities established across Ireland and Britain, we sought to comparatively profile the demographic histories of these genetic communities across different timescales by separating IBD segments into bins according to their lengths. We further used these bins to estimate temporal migration rates in Ireland and the UK, and the ancestral contributions from other European populations over three time periods. Since the time frames of IBD segment bins have wide ranges^34,39^, we refer to the bins by their length noting their age relative to each other, i.e. more recent, older etc. We complemented these methods with estimating levels of Runs-of-Homozygosity (ROH) for signals of inbreeding and/or isolation (see Methods). Lastly, we estimated changes in the effective population (N_e_) sizes over the past 100 generations using IBD segment data (See Supplemental Note 2).

#### Changes in structure across time

Comparing IBD sharing profiles in Ireland and Britain, we find evidence of shifting demographic relationships over time. We observe that approximately 100 generations ago (IBD length bin [1,3cM)) Irish communities on average shared more and longer IBD segments than most English and Scottish communities. (Fig. 3a and Supplemental Table 4). Further, the variation of sharing patterns within the Irish communities is subtler in comparison to their British counterparts, suggesting greater homogeneity (Fig. 3a-c and Supplemental Table 4). Examining patterns of IBD sharing between the Irish and British communities reveals change across more recent IBD length bins, demonstrating subtle shifts in population structure. We observed that elevated shared IBD levels between Irish communities (Supplemental Fig. 4a-c and Supplemental Fig. 5a-c) is primarily due to sharing within the [1,3cM) bin and it gradually decreases as the IBD bin lengths increase. This indicates an older signal of relative isolation in the Irish communities when compared to their general British counterparts. This corroborates our F_ST_ findings where the Irish communities have a lower median F_ST_ of 2.20 × 10^-4^ compared to UK communities (F_st_=9.51 × 10^-4^; Supplemental Note 1 and Supplemental Table 5).

**Fig. 3.**
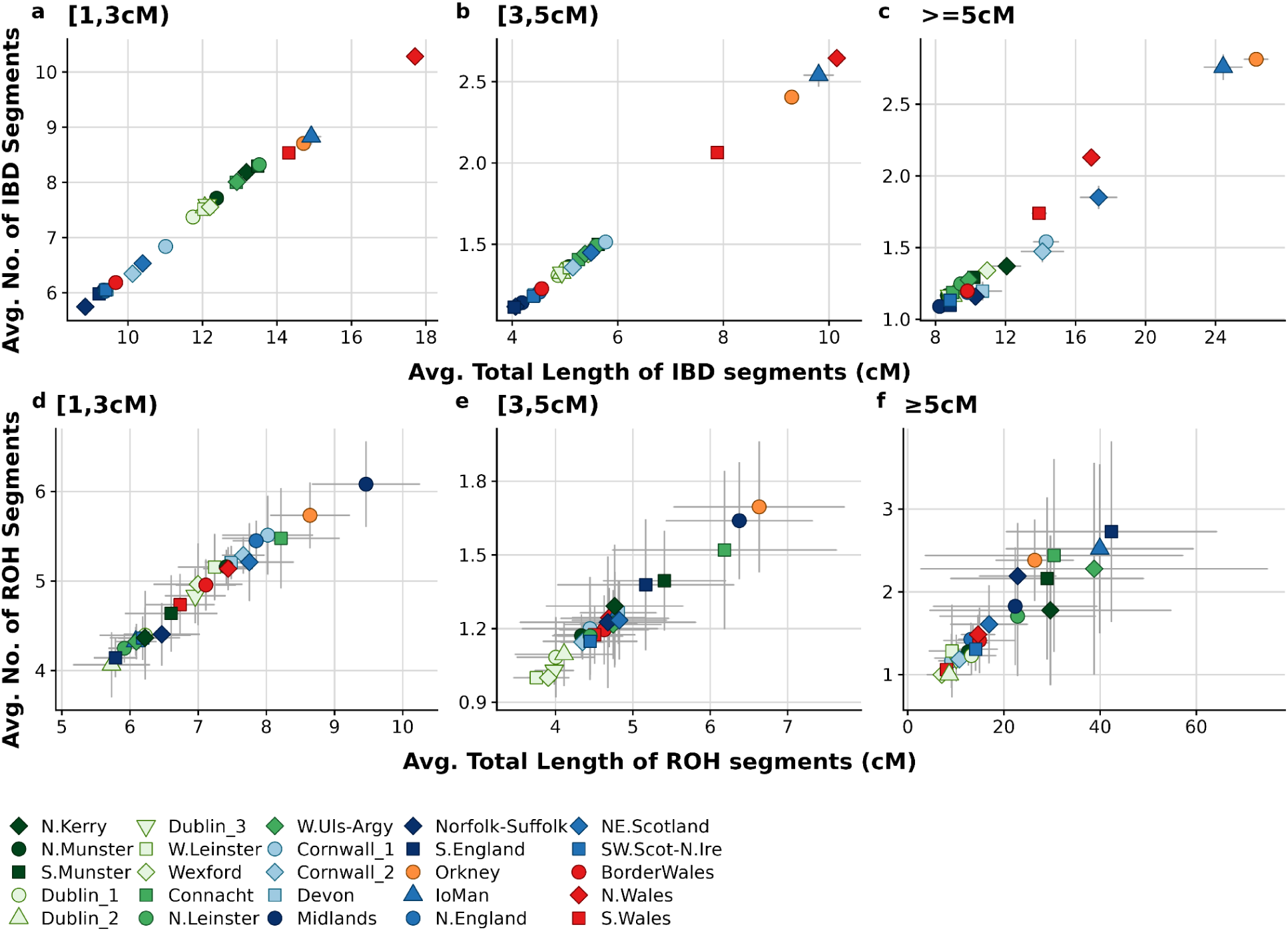
Inferring demographic history from the extent of haplotype and ROH sharing. (a-c) The scatter plots show the extent of IBD shared within every 3rd level genetic community after the IBD segments are segregated based on their length. Each point represents a 3rd level genetic community. The x-axis shows the average total length of IBD segments shared between pairs of individuals within the genetic community while the y-axis indicates the average number of IBD segments shared between pairs of individuals. (d-f) The scatter plots show the extent of ROH shared within every 3rd level genetic community after segregation based on their length. We plotted the average number of ROH segments shared between pairs of individuals in the community and the average total length of ROH segments shared between pairs of individuals to identify population isolates. The grey lines projecting from each point is the standard error.

In the [1,3cM) IBD length bin, we observed that IBD sharing increases as we move from the east to the west of the island (Fig. 2a), signalling greater isolation in the west of Ireland ∼100 generations ago. The within community sharing in Ireland however, remains significantly lower than highly isolated communities such as Orkney, N. and S. Wales and the Isle of Man across all length bins (Fig 3a-c). Irish communities have a similarly high degree of haplotype sharing as the Scottish, Manx, Welsh, and Orcadian groups in the [1,3cM) IBD bin (Supplemental Fig. 4a & 5a). However, while the Irish show equivalent values with the Manx and SW.Scottish groups in the [3,5cM), it diminishes with the NE.Scottish and Orcadian genetic communities in IBD bins ≥3 cM. In contrast, the N.Welsh genetic community appears to share slightly more IBD segments in the [3,5cM) and ≥5 cM bins (Supplemental Fig. 3a-c & 4a-c & 5a-c), as do the Cornish communities. These changing affinities between populations across time likely reflect complex demographic relationships which may be better understood by considering migration rates and isolation across time.

Population size and isolation can leave detectable signals on the distribution of ROH segments in a community. ROH segments in the [1,3cM) length bin indicate signals of isolation in the English Midlands, Orkney, Cornwall, and Connacht genetic communities 100 generations ago (Fig. 3d). This signal persists in the [3,5cM) ROH length bin for the Midlands, Orkney and Connacht genetic communities. However, the distribution ROH segments indicate recent isolation in the S.English, Manx, and W.Ulster-Argyll groups in >5cM length bin. Overall, ROH shared within the Irish, Scottish and English groups are comparable (Supplemental Table 6), the Orcadian, Manx and Welsh groups share higher levels of ROH. The SNP-based inbreeding coefficients (F_is_) and ROH-based coefficients (F_ROH_) for the populations indicate that the patterns of ROH and IBD-sharing observed is more likely due to small effective population sizes rather than consanguinity^40^ (Supplemental Fig. 8).

#### Changes in population size, isolation and migration

Haplotype sharing patterns within Ireland and Britain can provide insight into the sizes and movements of populations, providing context for observed genetic structure. Irish communities show relatively high IBD-sharing with little variation in sharing between Irish communities (Supplemental Fig. 4) indicative of low effective population sizes (Fig. 4). Using IBDNe, we observe that this relatively homogenous sharing pattern is reflected in the similar N*_e_* estimates of the Irish communities across the island 100 generations ago (Supplemental Fig. 7a-c, Fig. 4). Two-thirds of the Irish communities show a reduction in N*_e_* 40 generations ago, specifically the Wexford, N.Leinster, and S.Munster genetic communities which show 38-72% reduction in population size between 100 and 40 generations ago (Supplemental Table 7). We see a further reduction of 10% in the effective population size in the Wexford genetic community while there is an exponential increase (18-100%) in N*_e_* in the other Irish genetic communities (Supplemental Table 7). In addition, the N.Kerry community appears to have had a population expansion and followed by contraction within the past 30 generations (Supplemental Fig. 6a) which could be due to its small membership or may reflect cryptic relatedness just above our threshold of relatedness filtering (see Methods).

**Fig. 4.**
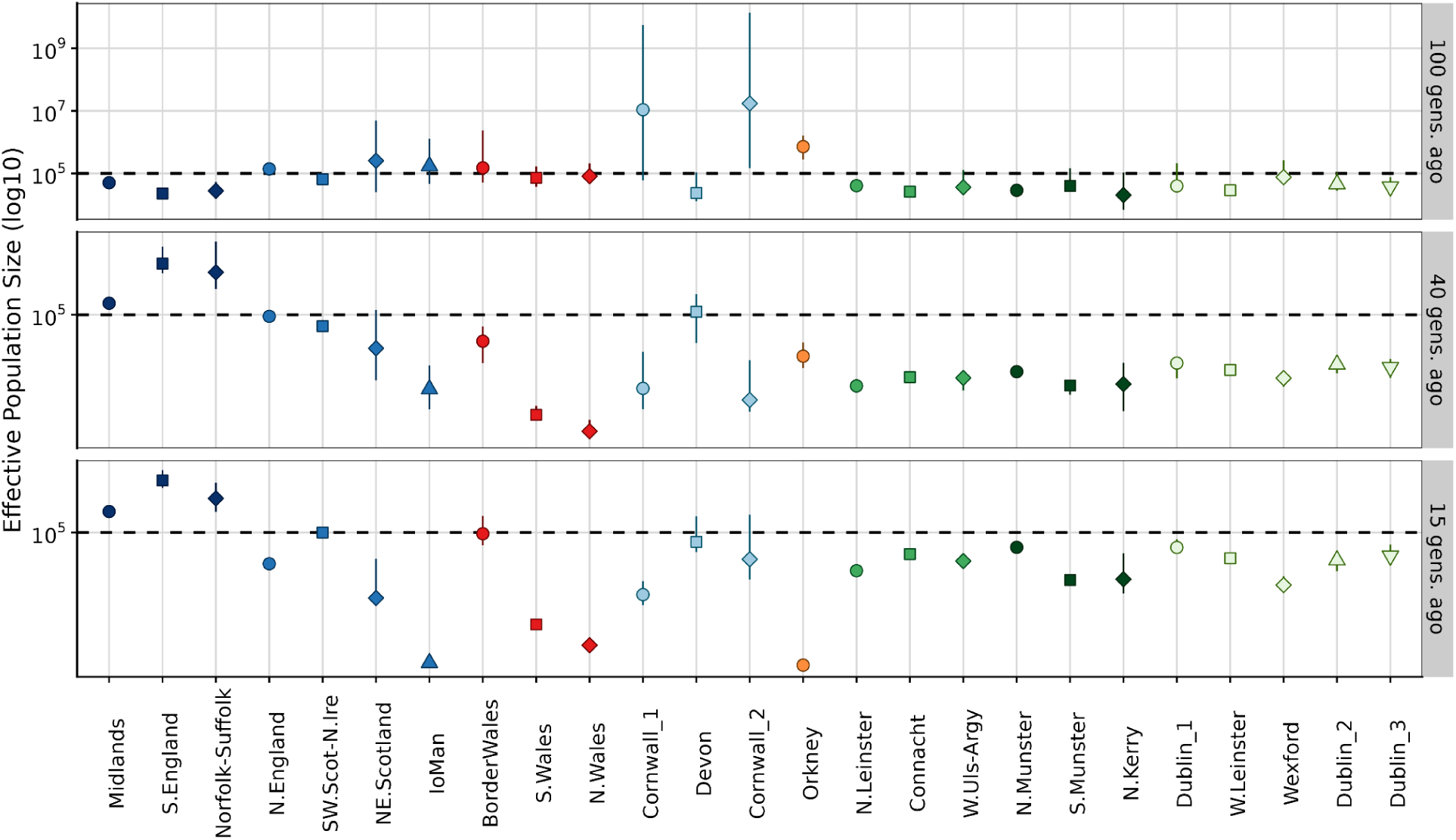
Effective population size (N*_e_*) estimates of communities in Ireland and Britain. The dot plots show snapshots of effective population size estimates of the genetic communities in Ireland (shades of green) and Britain (shades of blues, red, and orange) corresponding to the IBD length bins [1,3cM), [3,5cM) and ≥5cM. The third-level genetic communities in Ireland and Britain are represented along the x-axis and are indicated by the shape and colour of each point. The y-axis represents the effective population size estimates (N*_e_*) of the third-level genetic communities and is in log_10_ scale. The coloured lines from each point represent the 95% confidence intervals of each N*_e_* estimate.

Within the UK, there are elevated levels of haplotype sharing across varying bins of IBD or ROH length. The Orcadian, Manx, and the N. and S.Welsh genetic communities (and to a lesser extent Cornwall) demonstrate features of isolation (Fig. 2a-c and Supplemental Table 4). For example, this is reflected in their low effective population sizes over time, especially N.Wales, which are consistently lower than other British communities and show evidence of recent population contraction (Fig. 4, Supplemental Fig. 7d-g and Supplemental Table S7). By performing a PCA on an IBD-sharing matrix from IBD-segments in bins [1,3cM) and [3,5cM), principal components 1 and 2 resolved these communities from the Irish and other British communities (Supplemental Figure 6a-c). PCA on the average total length of IBD shared also shows that the Scottish, Manx and N.Irish groups are on a west-to-east cline between the Irish communities and British communities. The Cornish communities appear to be more recently isolated (Supplemental Fig. 6d) with their population sizes exponentially decreasing over the course of 80 generations, matching an enrichment of within-community IBD sharing >5cM. These Cornish communities share marginally more IBD with the Devon community and then the S.English community when compared to the other British communities (Supplemental Fig. 4 & 5). In contrast, the S.English communities see a steady increase in N*_e_*over time (Supplemental Fig. 7e), though with a common contraction in N_e_ followed by a recovery around 10 generations ago that is observed in nearly all clusters.

The observations of isolation in periphery communities above are reinforced by the migration rate surfaces (Fig. 5). We observed that the Orkney islands, Isle of Man, Wales, Cornwall and Devon show low migration rates to and from the mainland both in the older [1,3cM) and more recent (≥5cM) IBD bins, further supporting their isolation. The effective population size dips consistently over time in NE.Scotland, Isle of Man, Orkney Islands and Cornwall (Supplemental Fig. 6). There are further stable migrational barriers between north and south Wales, reflecting long standing structure within Wales. Additionally, there was little migration between the Scottish Lowlands and Highlands and Britain in the older [1,3cM) IBD bin. However, we observed a new migration corridor opening between the Scottish Lowlands and N.England in the more recent IBD bin (>= 5cM), while the Highlands remain isolated from the rest of the island (Fig. 5a-b). Interestingly, and supportive of analysis of IBD sharing across all the IBD length bins, N.Wales shows a historically smaller N_e_ around generations 30-50 than Orkney or the Isle of Man.

**Fig. 5.**
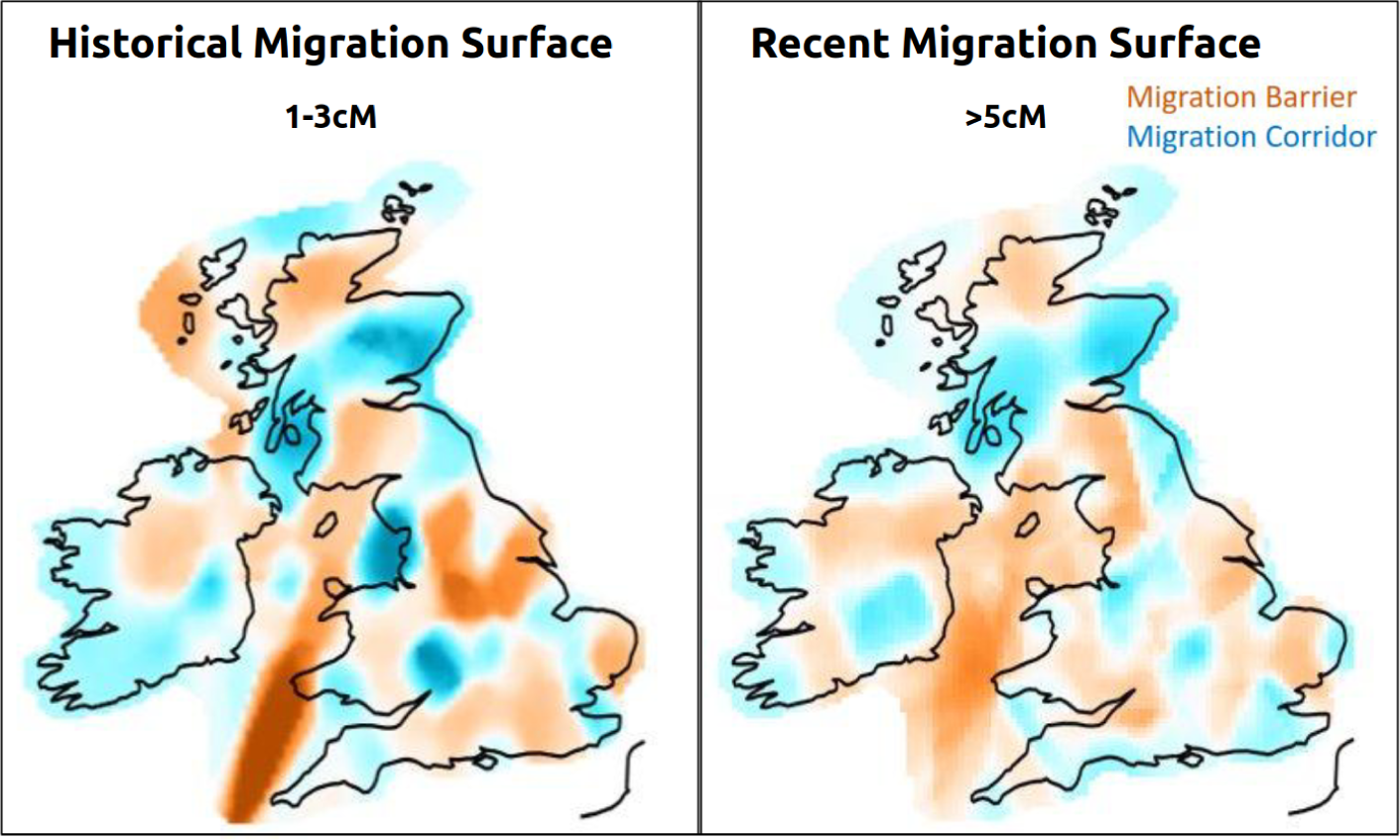
Temporal migration rate surfaces in Ireland and Britain. The plots show the estimated migration rate surfaces in Ireland and Britain in two time periods. The barriers are coloured in red while the migration corridors are coloured in blue. The outlines of the countries were sourced from Global Administrative Areas^59^. The figure was produced using the `plot_maps` function in the plotmaps package in R (ver. 4.2.1).

To further contextualise the structure and signals of isolation in Ireland, the migration rate surfaces (Fig. 5) show that the only stable migration corridor that consistently existed between Ireland and Britain is between northeast Ireland and southwest Scotland, demonstrating the isolation of the island. The within island migrational cold spots however shifted across time from central-West Ireland in the [1-3cM) IBD bin to isolating Leinster from the rest of the island 15 generations ago (Fig. 5), separating Kerry from Galway, which may explain the signals of isolation in Connacht. Additionally, in the north of the country, a shift in migration corridors links the west and east of Ulster, likely reflecting gene flow from Scotland across Ulster.

#### Continental European ancestry in Irish and British communities

We also looked into the European ancestry contributions to our genetic communities across time using PCA of European IBD sharing (Fig. 6 and Supplemental Table S8) to complement the previous results. In the oldest [1,3cM) IBD bin, we observe that English genetic groups separate out from Irish communities on axes driven by Germanic-Swedish-Norwegian ancestries. Scottish/Northern Irish communities demonstrate more Swedish influence while there appears to be strong North and North-Western Norwegian influence in the Manx and Orcadian groups (Fig. 6a and Supplemental Fig.9-12a-c). In the recent-history bin (>5cM) however, there appears to be more of a West-Germany ancestry signal in the English communities while there is more of a Swedish-Finnish-Norwegian component in the Scottish groups. The Norwegian signal persists in the Manx and Orcadian groups (Fig. 6b and Supplemental Fig.9-12a-c).

**Fig. 6.**
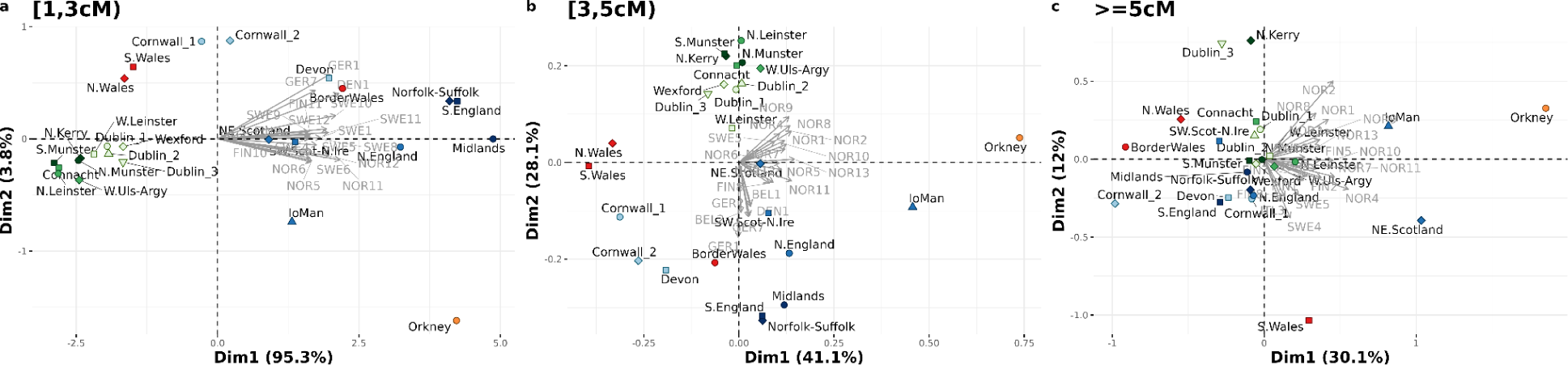
Biplots of regional European ancestry in Irish and British groups. We subset the dataset into IBD length bins and performed PCA on the average total length of IBD shared between the regional European genetic clusters and the Irish and British genetic communities. In these figures, we show the top 2 principal components (Dim1 and Dim2) and the PC loadings of individual variables (the regional European clusters) in grey that indicate how their influence changed over time periods.

Focussing on the contributions of European ancestry to Ireland over time, we find a strong signal from north Norway and western France and to a lesser extent from Sweden in the Irish communities in the [1-3cM) IBD bin (Supplemental Figs. 9-12). There appears to be a more recent contribution of ancestry in the [3,5cM) IBD bin from North-North-West Norway and Sweden in Ireland, specifically in the W.Leinster community (Supplemental Figs. 9-12).

## DISCUSSION

In our study, we used patterns of IBD-shared between 6,574 individuals with regional ancestry to infer recent demographic histories across Ireland and Britain — providing context for genetic communities described here and in literature. We leverage IBD segments to show that genetic structure and regional genetic difference in Ireland is subtler when compared to Britain. We also show temporal changes in regional migration rates and population sizes in Ireland and Britain, demonstrating signals of population isolation in peripheral genetic communities (e.g. Wales, Isle of Man and Orkney) and growth in others (e.g. South England). Additionally, we provide genealogical context to the Irish genetic communities through surname analysis.

In agreement with previous studies^1–3,21^, we found the genetic communities detected largely divided into Irish and British-like genotypes with sub-communities that resolved along provincial/historical kingdom boundaries in Ireland and country/administrative boundaries in the UK. Within Ireland, we identified novel genetic communities with geographic provenance in north Kerry and north Leinster within Ireland, whilst within the UK, we detected novel communities from the Isle of Man and regions within England (East Anglia, south England and the Midlands, and east and west Cornwall). Using the geographic provenance in our dataset we were able to predict regional Irish or British ancestry within larger datasets such as the UK Biobank, demonstrating the utility of well-characterised reference datasets to provide greater value in studying regional health genomics in large biobanks.

With the genetic structure of Ireland and Britain as the basis, we leveraged IBD segments and ROH to infer over time and in relation to the identified communities, changes in (a) effective population sizes, (b) the extent of genetic relatedness between them, and (c) relatedness between them and regional European genetic clusters. We showed time-sensitive changes in the demographic profile of different regions across Ireland and Britain by utilising IBD-segments across different length bins. Whilst the confidence intervals of the generational estimates around these bins are wide, we are able to show observable changes across different regions. The relative isolation of Irish groups was demonstrated by the slightly elevated levels of IBD shared between them regardless of the IBD bin lengths and by its smaller effective population size compared to the UK in general. We also found features of isolation in British regions such as the Isle of Man which shows a signal of isolation similar to Orkney, although smaller in magnitude. Interestingly, we observed elevated levels of IBD sharing between the Irish and Scottish, Welsh, Manx and Orcadian groups in the smaller IBD bins that disappeared in the ≥5cM IBD bin, suggesting a shared historical Celtic/Norse-Viking signal in these populations. The affinity of the Manx group in our dataset to the Irish population is in direct contrast to the report from Gilbert et al in 2019 which used a different cohort of Manx individuals. The report showed a high proportion of English ancestry in Manx individuals. The observed difference between our observation and that reported by Gilbert et al could be due to different ascertainment schemes from the Isle of Man. Additionally, our time-sensitive IBD-bin method is able to highlight novel results in Wales. We found an excess of within-group sharing in smaller IBD-length bins which disappears in more recent bins, suggesting the primary source of genetic isolation observed in Wales is older than in Orkney or the Isle of Man. By inferring regional demographic histories, we confirmed the Orcadian genetic community has historically been genetically isolated for the past 100 generations. We also observed a consistent gene flow between the Orcadian community and the northeast Scottish group.

By estimating contributions of European ancestries in the Irish and British genetic communities across timescales, we found evidence of gene flow between Scandinavia and the English, Scottish and Irish groups in the smaller IBD bins but across all IBD bins within the Orcadian and Manx communities. We also detected an affinity between modern populations in northwest Germany and the English communities in the smaller IBD bins, consistent with the influence of Germanic migrants in the Migration Period^5^ in the region^21^. Further, by separating out signals of sharing with European groups, extending previous findings using haplotype “painting” of chromosomes, we shed light on the changing landscape of European affinity across the islands. This particular analysis touches again upon the question of the genetic legacy of Norse Viking activity on the modern Irish genetic landscape. We observed a higher similarity between Irish genetic communities and modern Norwegian haplotypes in smaller IBD segment bins that is concurrent to the signal in Orkney and the Isle of Man and consistent with a Norse-Viking origin. Recent work with ancient DNA in Scandinavia has demonstrated significant gene flow from Ireland and the British Isles into Viking-era Scandinavia^9,10^ including the regions of Norway previously shown to have an excess sharing in Ireland^3,8^. Our results here are consistent with an inflation of present-day Norwegian sharing to present day Ireland due to this historical Irish gene flow to Viking-era Norway.

However, the uneven sampling participants across Ireland and the UK has led to overrepresentation within some of the identified genetic communities. Samples from Scotland were predominantly from the North-East and South-West of the country, preventing analysis of known Scottish genetic clusters^8^ in between. Generally, the British sampling was more rural which limited analysis of large urban areas. Further, we had few samples from north-west and south-west of Ireland, although this would match population densities across the island.

In conclusion, we show that the regional genetic differences in Ireland are subtle and can largely be attributed to older genetic sharing whereas the observed genetic differences in Britain are largely along administrative boundaries. Through analysis of haplotype sharing, we identified specific regional Irish and British genetic communities that have been genetically isolated over different time periods, and highlight the signals of contracted population size in Wales. Therefore, our results from studying the demographic histories of regional Irish and British populations and their changing affinities over time motivate the study of rare variants within these populations, to determine if they indeed stratify by patterns that match the demographic history of the population. Additionally, we have shown that these results can be used to predict regional ancestry of Irish participants in larger datasets such as the UK Biobank.

## MATERIALS AND METHODS

### Datasets

Genotype data from individuals of British and Irish ancestry was accessed from four studies: the Irish DNA Atlas (*n* = 198) (Atlas)^2^, the Trinity Student Study (*n* = 2,232) (TSS)^41^, the Trinity Irish ALS case-control cohort (*n* = 991) (ALSCCC)^42^ and an updated version of the People of the British Isles dataset (PoBI)^21^ with additional genotypes (*n* = 6,294). In addition, we accessed European reference haplotypes (*n* = 6,916) from the WTCCC2 Multiple Sclerosis dataset^43^(WTCCC2-MS) to estimate temporal differences in ancestry proportions. We subset the UK Biobank genotypes^38^ to participants who reported their place of birth as “Ireland’’ or “Northern Ireland’’ (F.21000) and they mapped with the Irish references on the principal components (F.22009) plot (*n* = 7,575).

Geographical provenance was available for two datasets: the Irish DNA Atlas and the PoBI datasets. The great-grandparents of all individuals recruited for the Irish DNA Atlas were born within 50 km of each other^2^. As for the PoBI dataset, the grandparents of all individuals in the study were born within 80 km of each other^21^. We assigned geographic provenance to the participants with the average coordinates of birthplaces of the great-grandparents or each participant for Irish DNA Atlas and those of the grandparents for PoBI. The geographic data for the place of residence was available for approximately half of the individuals in the ALSCCC dataset.

We obtained the necessary permissions to work with the datasets used in our study. Please refer to the original papers for the ethics statements^2,3,21,38,42^.

### Quality Control

Using PLINK (ver. 1.9)^44^, we first processed individual datasets by excluding any strand-ambiguous SNPs and included autosomal SNPs. For the Irish-British analysis, we combined the Atlas, PoBI, TSS, and ALSCCC datasets. For the European ancestry analysis, we combined the Atlas, PoBI, TSS, and ALSCCC datasets with the WTCCC2-MS dataset.

Datasets were merged using the PLINK command --merge. We removed SNPs with minor allele frequency (MAF) <2%, missingness >5%, and a Hardy-Weinberg Equilibrium (HWE) p-value <1e-6. We excluded individuals with SNP missingness >5% and excluded SNPs in regions known to be in linkage disequilibrium^45^. Pairs of related individuals (up to the 3rd degree) were identified using KING (ver. 2.2)^46^ and one random individual from each pair was removed. To calculate an unlinked PCA, we utilised a set of SNPs that were pruned with respect to linkage disequilibrium using the PLINK command --indep-pairwise 1000 50 0.2, performing PCA using the --pca PLINK command on samples for the Irish-British and the European ancestry analyses.

The final Irish-British dataset had 6,574 samples and 298,297 markers while the European dataset had 13,029 samples and 194,714 markers. This level of coverage has been found to be sufficient when considering longer (i.e., > 3cM) segments of IBD or ROH^47^. The geographic data for the Trinity ALS dataset has been jittered to preserve anonymity. The combined Irish-British dataset and the Irish subset from UK Biobank had 12,690 samples over 69,788 markers.

### Phasing and IBD Segment Detection

Unphased genotypes were converted to VCF files using PLINK and separated by autosome. Using SHAPEIT4^48^, we phased the samples with default settings and the GRCh37 build genetic map for recombination distances generated from HapMap. IBD segments ≥1cM in length were detected using refinedIBD^27^ and adjacent IBD fragments within 50kbp and 0.06cM of each other were merged using the merge-ibd Java script — merging putative IBD segments which were broken by either phase or genotyping errors. To extract data reflective of different time periods, autosomal IBD data was merged and subsets were created based on length of IBD segments. The extent of IBD shared between pairs of individuals was estimated by calculating the average total length of IBD shared and the total number of IBD segments^49^.

### Population Structure Detection

Using the summarised IBD data, we first constructed a network graph using every individual as a node and the average total autosomal IBD length shared between a pair of individuals as the edge (*igraph* package in R^50^). We exclude outlier connections of oversharing to account for cryptic relatedness, excluding connections with a sum of all segments ≤ 99.995 percentile of IBD length - 122.87 Mbp. The Leiden community detection algorithm^35^ was then applied on this network. We used the *rleiden.community* function within the *leidenAlg* package in R^51^ with a max.depth of 3 (i.e. 3 recursions of the clustering process) and a *min.community.size* of 100 (i.e. communities of 100 individuals were not divided in a subsequent recursion). The communities were labelled reflective of their geographic membership.

To estimate reproducibility of these Leiden clusters, we ran the leiden.community function with the same settings for 100 iterations. Using the genetic communities from the first iteration as the basis, we calculated the proportion of individuals who were grouped together in the same genetic community at the three levels of recursion for every subsequent iteration, and then averaged across the original clusters.

To investigate the hierarchical structure of the genetic communities, we computed Euclidean distances between the groups using the haplotype-sharing matrix generated by pbwt-paint (https://github.com/richarddurbin/pbwt/blob/master/pbwtPaint.com) between pairs of individuals in the dataset to generate a distance matrix using the dist function in R. The *hclust* function was applied on this distance matrix to generate the final dendrogram. We then validated the dendrogram using the pvclust function from the pvclust package with 10,000 bootstrap resampling. We estimated genetic distances between communities using F_ST_ as caculated in the admixtools package in R^50^, and measured cluster robustness with TVD estimates^21^.

### Predicting Regional Ancestry within the UK Biobank

We used PLINK to perform principal component analysis after combining our Irish and British genotypes with the UKB Irish subset. We then randomly split the Irish-British reference data 7:3 to get training and validation datasets. We scaled the first five principal components of the training set and trained a Naive Bayes classifier with the data and their corresponding second-level genetic community label with the default parameters (*naiveBayes* function in the *e1071*^53^ R package). To evaluate the accuracy of the model, we predicted the second-level genetic community label of the validation dataset and calculated the concordance between the observed and the predicted label. We predicted the second-level genetic community labels for the Irish subset of UKB using the model.

### Estimating Extent of Genetic Relatedness

For every genetic community, we first created subsets of IBD segment sharing data based on the length of IBD segments: IBD segments of (a) between 1 to 3cM ([1,3cM)), (b) between 3 to 5cM ([3,5cM)) and (c) greater or equal to 5cM (5<=IBD). These approximately correspond to (a) 100 generations ago, (b) 40 generations ago, and (c) 15 generations ago respectively^39^. To characterise their histories, we then plotted the average number of IBD segments versus the average length of total IBD segments shared for the different IBD length bin sizes. We calculated the total length and number of IBD segments shared between pairs of individuals in the dataset. Then, we generated summary statistics for every pair of 3rd level genetic communities. We generated a heatmap each after calculating the z-scores of the average total length and number of IBD segments respectively. Additionally, we generated principal components using the average total length of IBD segments using the prcomp function in R.

To estimate consanguinity in the Irish and British genetic communities, we estimated ROH within every genetic community using --homozyg with a minimum SNP count of 50, a scanning SNP widow size of 50 SNPs, minimum length of 1500 kb, maximum internal gap of 1000 kb length, a maximum inverse density of 50kb/SNP, a maximum missing calls in scanning window hit of 5, and a maximum number of heterozygous hits in scanning window of 1. We additionally estimated the F_ST_ ^54^ statistic, and the F_ROH_ ^55^ and F_IS_ ^54^ statistics for every 2nd level and 3rd level genetic community.

### Estimating Effective Population Sizes

To estimate changes in effective population sizes over time, we used IBDNe^28^. We selected IBD segments ≥4cM shared between pairs of individuals within the 3^rd^ level recursion of Leiden genetic communities in Ireland and Britain. We then ran the IBDNe program^28^ on this dataset with *nboots*=100 to generate confidence intervals of the Ne estimates.

### Migration Surface Estimation

We first limited the IBD sharing summary dataset to pairs of individuals both of whom had associated geographic data. It was further subset by IBD length to determine changes in migration surfaces over time: [1,5cM) for migration rates from approximately 100 generations ago and ≥5cM for migration rates from approximately 15 generations ago. A square matrix that captured the number of IBD segments shared between pairs of individuals was created. With the IBD sharing matrix, a file with the geographic coordinates of every individual and shape file with the coordinates of the outline of the Irish and British borders as input, we ran Migration and Population Surface (MAPS)^34^ to estimate time-resolved dispersal rates and population densities. We set nDemes = 300, numMCMCIter = 1×10^6^, numBurnIter = 5×10^5^ and numThinIter = 100 over 5 parallel chains. If the log-likelihood values from all the chains didn’t converge to similar values, we used results from the chain with the highest log-likelihood value to initialise the next round of estimations with the same parameters until the log-likelihood values converged. The figures were generated using the *plotmaps* package^34^.

### Estimating Timeline of European Ancestry in Ireland and Britain

After combining the WTCCC2-MS dataset with the Irish-British dataset, we detected IBD segments and sorted the IBD segments into length bins of [1,3cM), [3,5cM) and ≥5cM. With the regional European reference ancestry labels^2^ generated using fineSTRUCTURE, we summarised the extent of IBD shared between each European reference ancestry population and the 3rd level recursion of genetic communities within Ireland and Britain within the IBD length bins. We then generated PCA biplots^56^ from this data to detect the temporal contributions of ancestry from the reference European populations. We also generated heatmaps after scaling the average total length and the number of IBD segments shared between pairs of populations.

### Surname Analysis

Through the Irish DNA Atlas study, we had access to the surnames of the eight great-grandparents of every participant. These surnames were classified into 10 groups based on their origin (Supplemental Table 8) by Irish genealogists (S.O’R and M.M). Those whose surnames were unknown/ambiguous were assigned into the “Unknown” category. We first assigned the genetic community label to each great-grandparent and then created a two-way frequency table of the number of surnames by their origin and the genetic group to which they belonged. Using this table as input, we used the *assoc* function in the *vcd* package^57,58^ (ver. 1.4-11) to identify which surnames are enriched or depleted within each genetic community that uses residuals from a Chi-square test to infer the results.

## ACKNOWLEDGMENTS

We would like to thank all the participants of the Irish DNA Atlas, the Trinity Student Study, the Trinity ALS case-control cohort, the People of the British Isles, This work was supported by funding from Science Foundation Ireland (SFI) through the Centre for Research Training in Genomics Data Science under grant number 18/CRT/6214 and through the SFI FutureNeuro Research Centre under grant number 6/RC/3948 and co-funded under the European Regional Development Fund and by FutureNeuro industry partners. This work was also further supported by Science Foundation Ireland (17/CDA/4737 and SFI 15/SPP/3244) and the MND Association (879-791 and 979-799).

