## Supplemental Notes for "A genetic perspective on the recent demographic history of Ireland and Britain"

#### Supplemental Note 1

The results from *pvclust* show that the Irish communities resolved along geographic boundaries. The branch consisting of South Munster and North Kerry was similar to the communities from Connacht, North Leinster and North-West Ireland. Interestingly, North Munster grouped together with communities from Wexford, West Leinster, and Dublin and not with other Munster communities. Within the UK sub-branch, we observed the Northern Irish communities group together with North England and North-East Scotland. The Welsh and Cornish communities are grouped in their own sub-branches respectively. England formed its own sub-branch, where the English community split into sub-communities with individuals with ancestry from East Anglia, the Midlands and South England grouped separately, and branched with individuals with ancestry from Border Wales and Devon. The certainty of these broad Irish and British branches of the tree were confirmed using 10,000 bootstrap resampling with the *pvclust* implementation of *hclust*.

This reproducibility analysis of genetic community detection showed that reproducibility decreases with every recursion of the Leiden community detection algorithm. The median reproducibility of the first recursion was 99.95%. For the second recursion, the median reproducibility was 87.52% and 74.47% for Irish and British communities respectively. However, reproducibility was drastically reduced for the third recursion: it ranged between 15.82 to 38.41% for the Irish communities and 25.09 to 78.01% for the British communities (Supplemental Table 1). We further tested robustness through FST distances between our British and Irish communities which demonstrated that the genetic differences between them are subtle with the greatest differentiation observed between the Orcadian and non-Orcadian communities ($F_{ST}$ = 0.00186), in agreement with previous reports[1–3,21]. Interestingly, the North Wales ($F_{ST}$ = 0.00175) and Isle of Man ($F_{ST}$ = 0.00165) communities also have relatively higher levels of separation compared to the other Irish and British genetic communities (Supplemental Table 2). Within Ireland, the North Kerry, South Munster, and Wexford communities differed the most genetically from the other Irish communities. Genetic communities from South Leinster were similar to communities from Connacht-Leinster. The English and the Cornish communities showed more subtle genetic differentiation compared to the Orcadian, Welsh and Cornish communities. Total-variation distance (TVD) measures confirm that the communities detected by the Leiden algorithm are indeed robust — the p-values are significantly lesser than 0.01 (Supplemental Table 3).

### Supplemental Note 2

Demographic histories of the regional genetic communities were inferred by analysing IBD and ROH segments in length bins. We estimated the expected segment age for each length bin using equation s19 from reference 35 (reproduced here), assuming a large effective population size[34].

$$\lim_{N \to \infty} E[T|\mu \leq l \leq \upsilon] = 75\left(\frac{1}{L_1} + \frac{1}{L_2}\right)$$

Where $T$ is coalescence time (generations), $l$ is segment length (base pairs), $\mu$ and $\upsilon$ are the upper and lower segment length bounds of the bin (base pairs) and $L_1$ and $L_2$ are the upper and lower bounds rescaled to centiMorgan.

We chose to split ROH and IBD segments into the corresponding bins (a) 1 to 3cM [1,3cM), (b) 3 to 5cM [3,5cM), and (c) greater than or equal to 5cM (≥5cM). Using the methodology above[34], these length bins correspond to 100 generations ago, 40 generations ago and 15 generations ago respectively. An important caveat is that the estimates have very wide distributions, as well as the aforementioned assumption of population size.
