## Supplemental Figures for "A genetic perspective on the recent demographic history of Ireland and Britain"

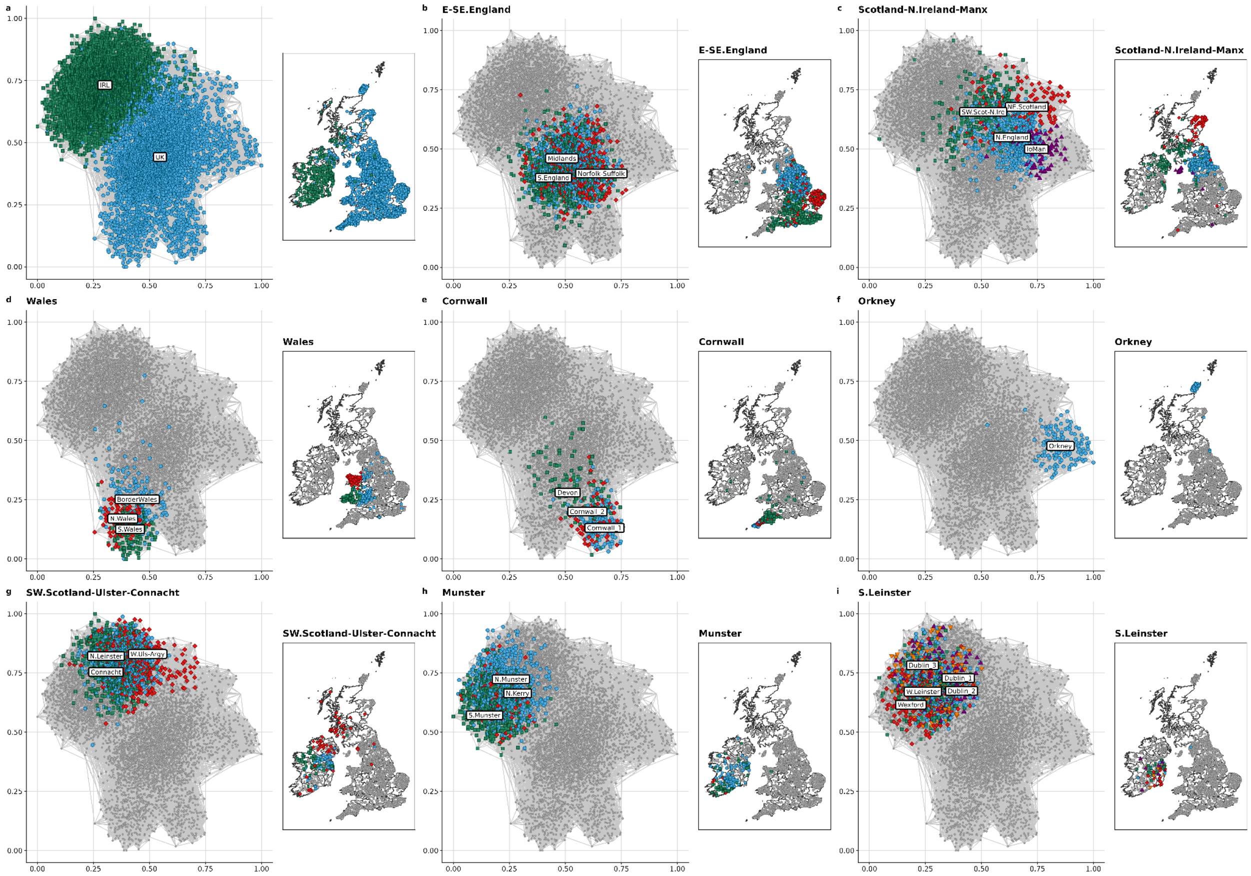


**Supplemental Fig. 1.** Third level genetic communities in Ireland and Britain

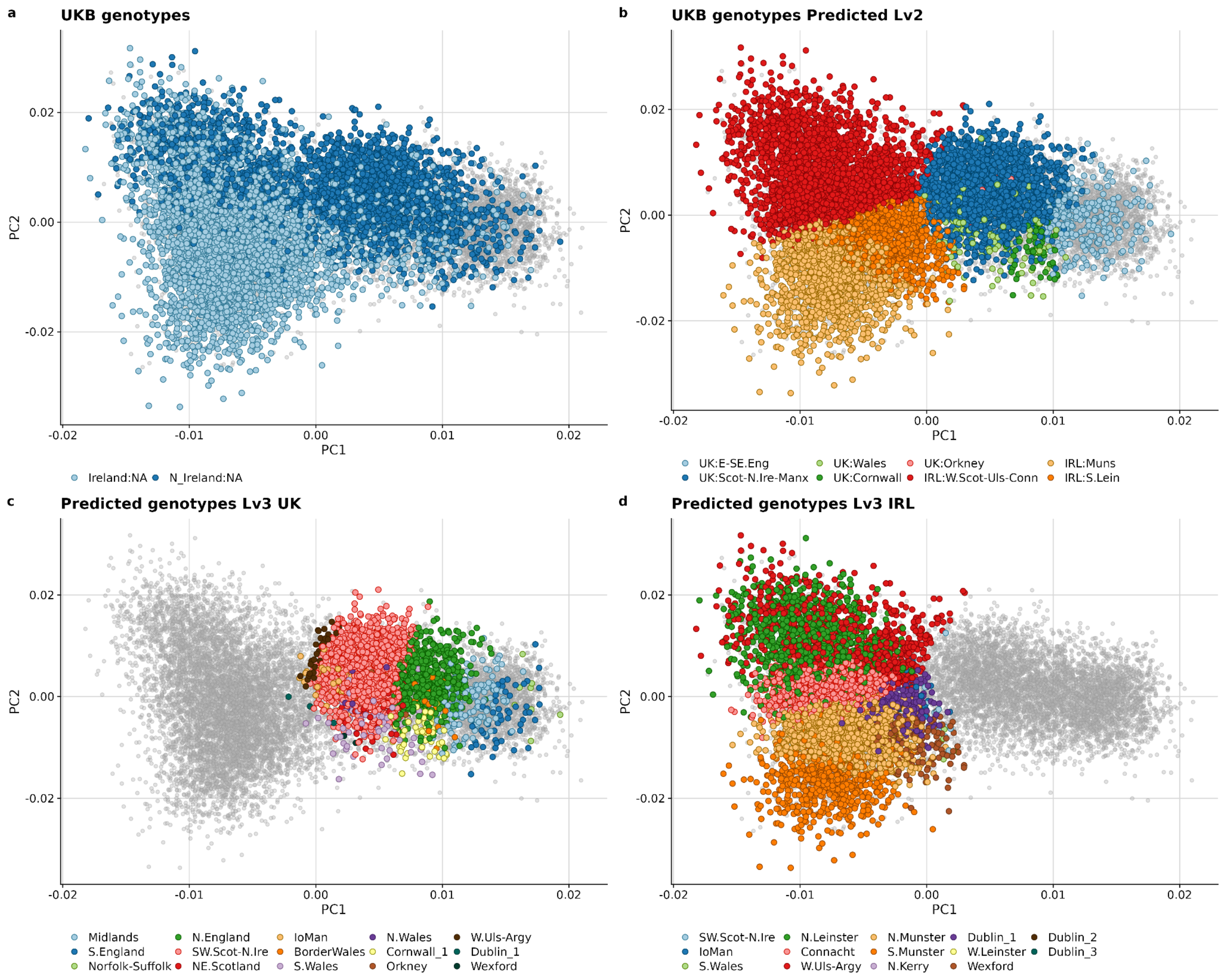


**Supplemental Fig. 2. Predicting the regional ancestry of the Irish subset of the UK Biobank participants.** (a-d) show the first two principal components generated using PLINK. Every point coloured grey is a participant from our dataset. (a) The two shades of blue denotes the reported place of birth of the participants within the UK Biobank. (b) The colours in this plot indicate the second level label assigned to the UK Biobank participant by our Naive Bayes model. (c &d) The colours in these plots indicate the third level label from UK and Ireland respectively assigned to the UK Biobank participant.

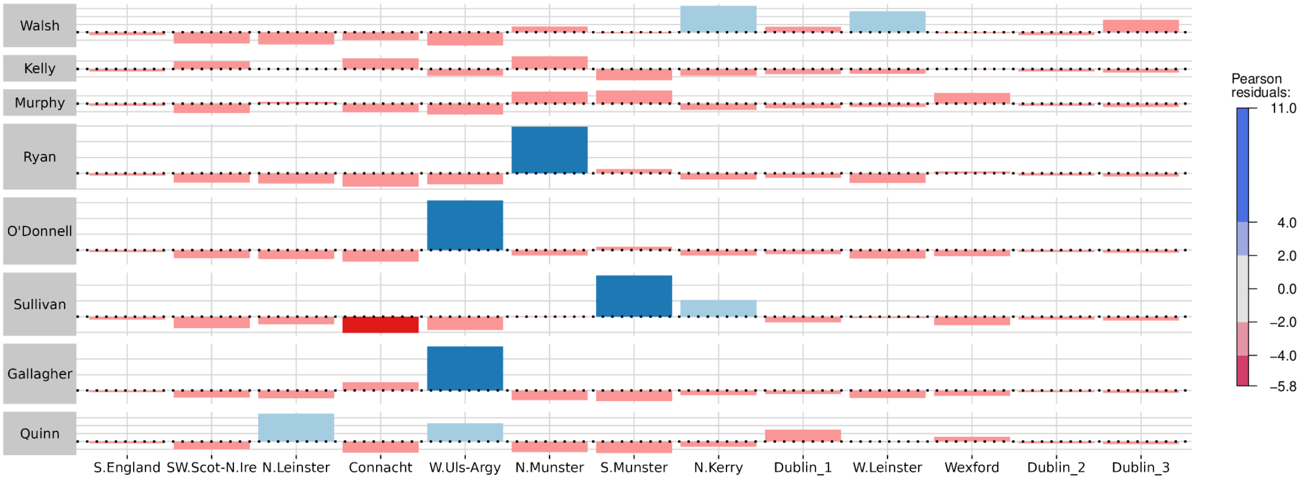


**Supplemental Fig. 3. Surname association with regions in Ireland.** Using a Chi-square test, we estimated the differential occurrence of surnames (taking the top 8 surnames in Ireland) within the Irish genetic communities. The colours of the bars indicate the Pearson residual value of the surname (along y-axis) within each genetic community (x-axis).

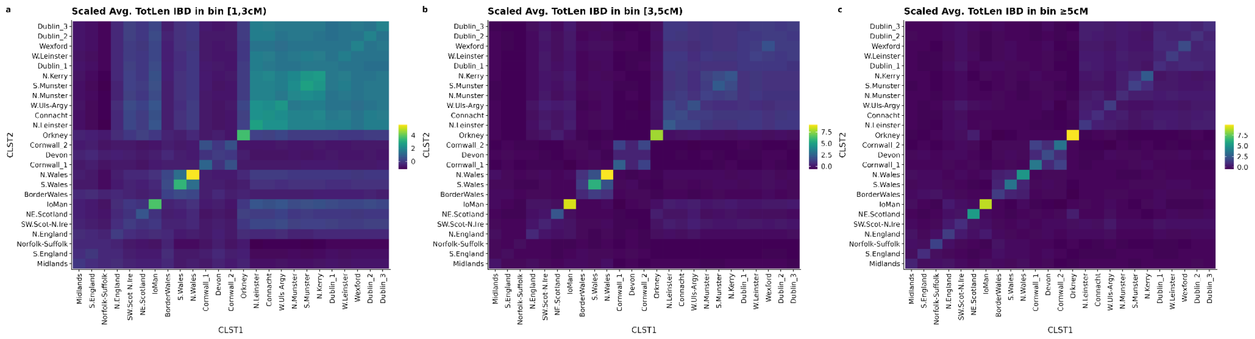


**Supplemental Fig. 4.** Heatmap showing the average total length of IBD shared between 3rd level communities in Ireland and the UK across length bins scaled (z-scores) by row

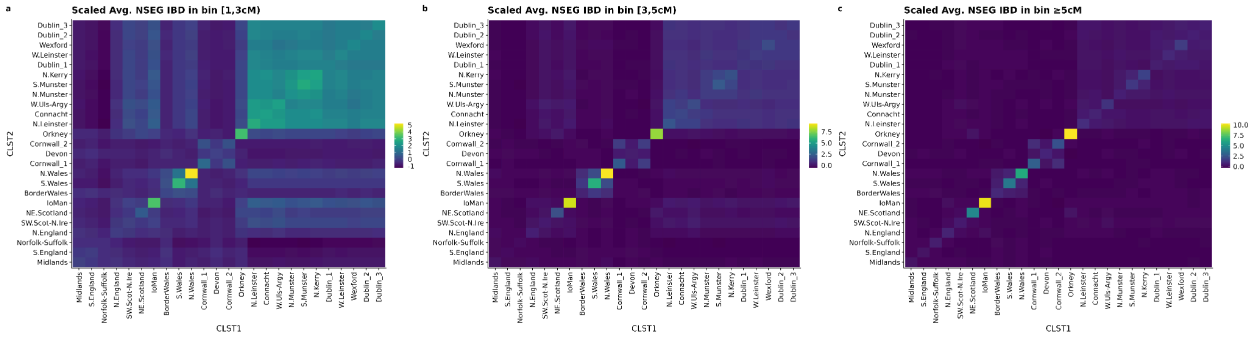

**Supplemental Fig. 5.** Heatmap showing the average total number of IBD segments shared between 3rd level communities in Ireland and the UK across length bins scaled (z-scores) by row

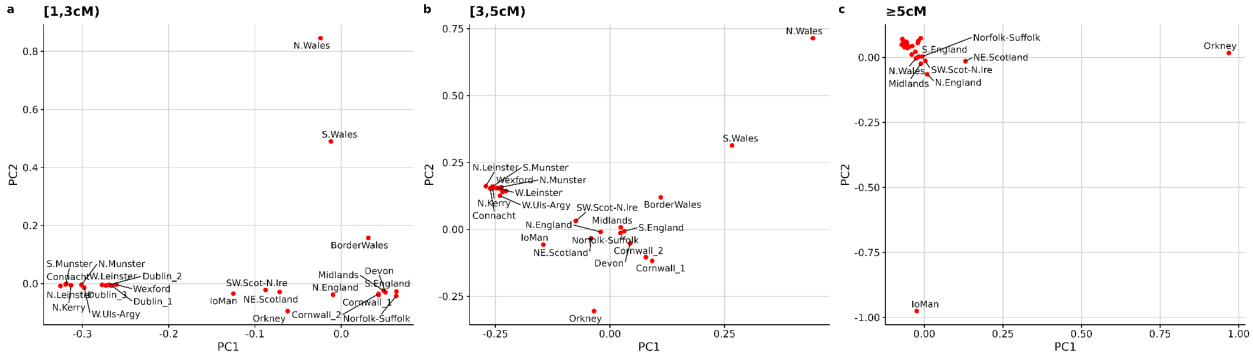

**Supplemental Fig. 6.** Principal components analysis on the average total length of IBD shared between 3rd level communities in Ireland and the UK across length bins. These figures show the top 2 principal components.

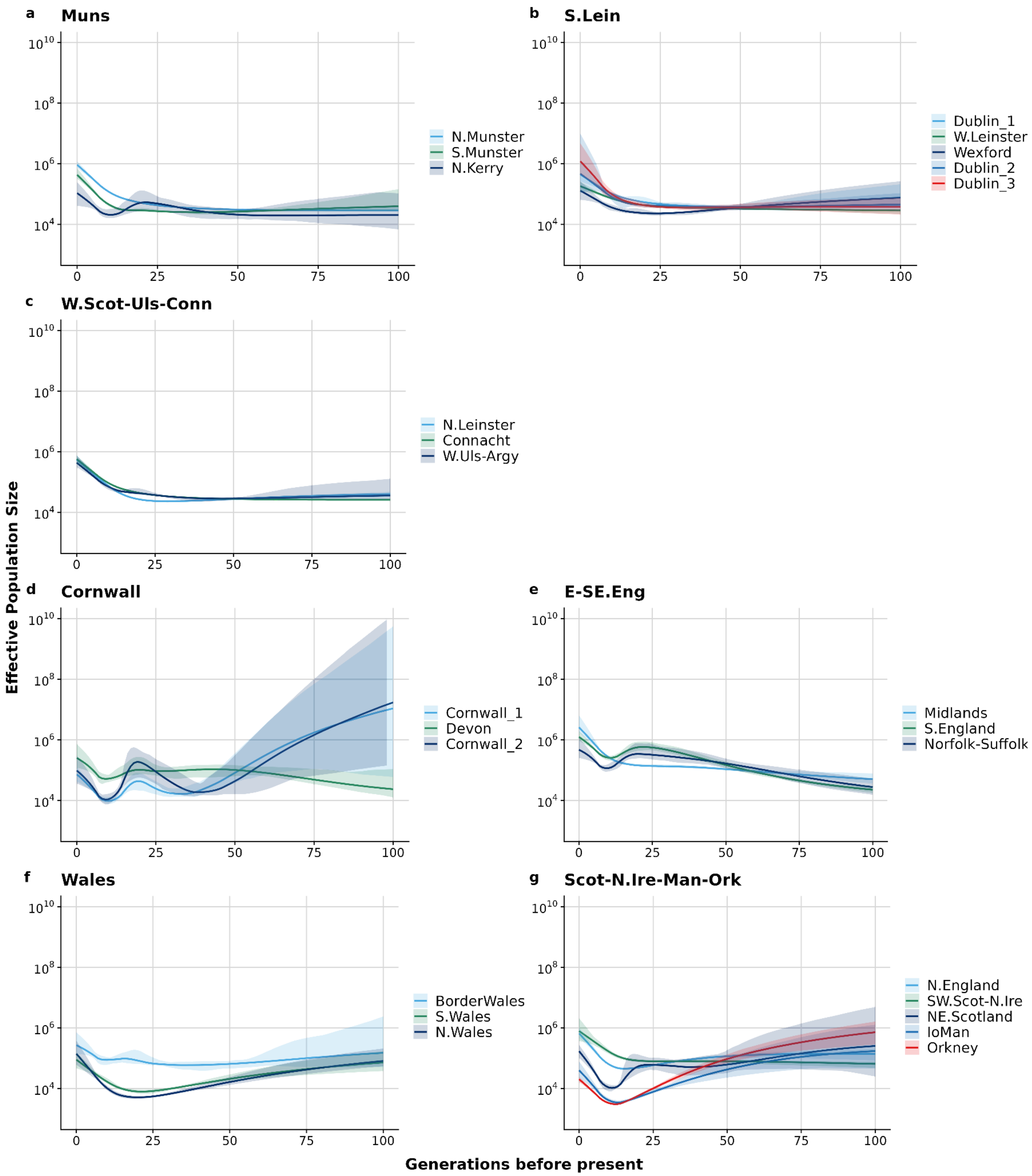


**Supplemental Fig. 7. Historical effective population size ($N_e$) estimates of communities in Ireland and Britain.** The line plots show the trajectories of genetic communities in (a-c) Ireland and (d-g) Britain. Each line represents a 3rd level genetic community and they are grouped by their parent/2nd level genetic community and the shaded region represents the 95% CI.

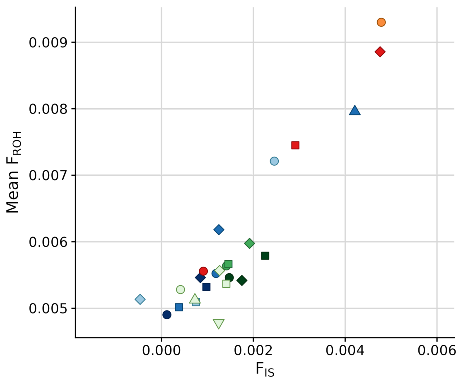


**Supplemental Fig. 8**. We tested for consanguinity by plotting Mean $F_{ROH}$ vs $F_{IS}$. The position of the Irish and British clusters along the y-axis is indicative of small effective population size. The colour of the points on the plots indicates the 2nd level genetic community while the shape of the points indicates the 3rd level genetic community.

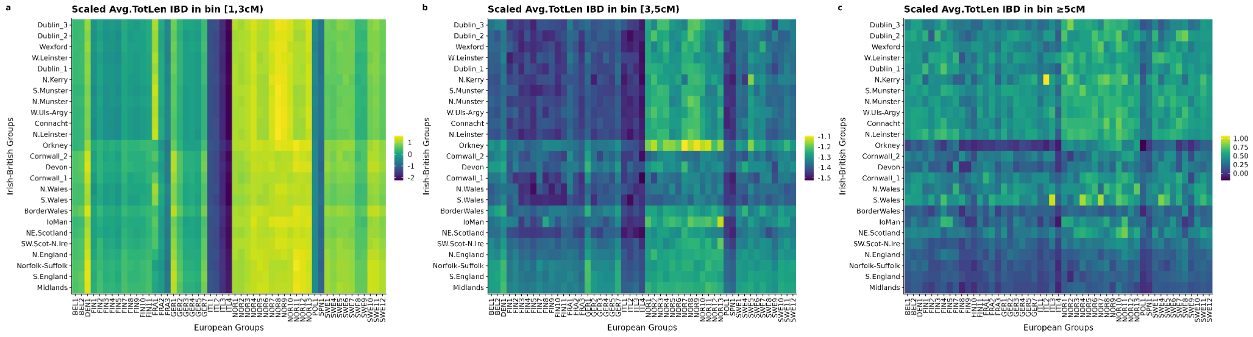


**Supplemental Fig. 9.** Heatmap showing the average total length of IBD shared between 3rd level communities in Ireland and the UK and regional European clusters across length bins scaled (z-scores) by row to capture variation in sharing profiles with European clusters within individual Irish and British genetic communities.

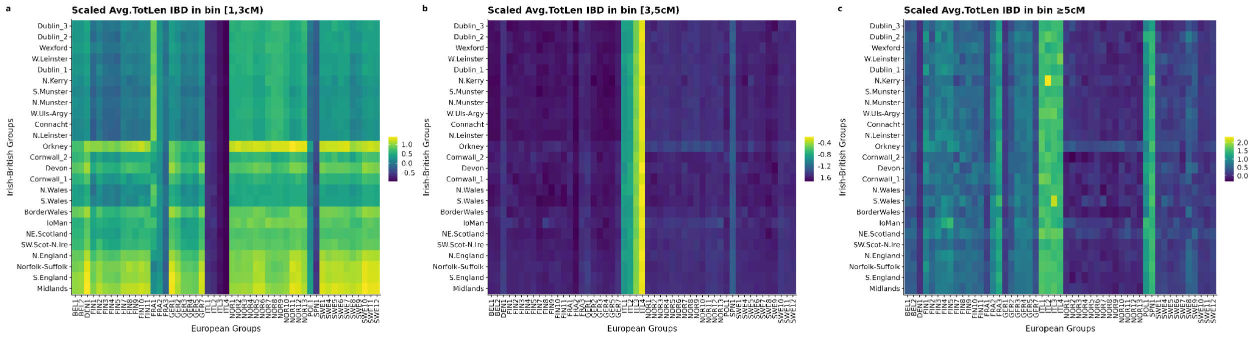

**Supplemental Fig. 10.** Heatmap showing the average total length of IBD shared between 3rd level communities in Ireland and the UK and regional European clusters across length bins scaled (z-scores) by column to capture variation in sharing profiles with European clusters across all Irish and British genetic communities.

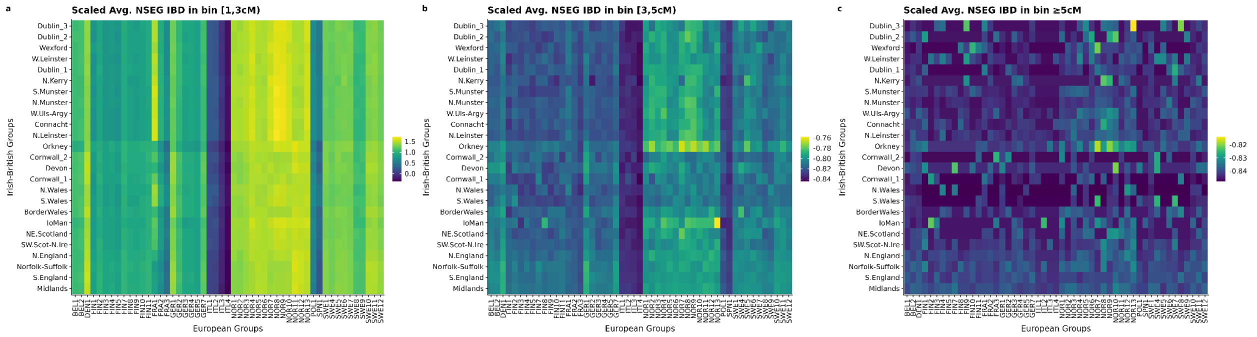


**Supplemental Fig. 11.** Heatmap showing the average total number of IBD segments shared between 3rd level communities in Ireland and the UK and regional European clusters across length bins scaled (z-scores) by row to capture variation in sharing profiles with European clusters within individual Irish and British genetic communities.

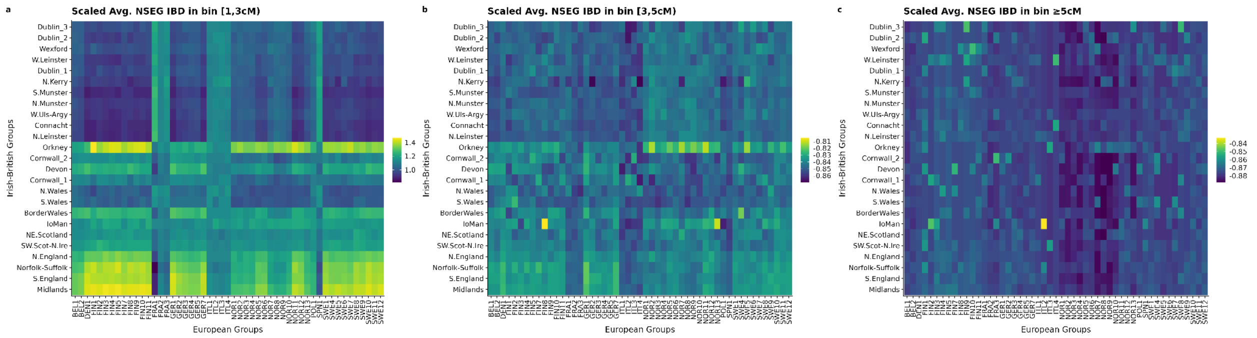


**Supplemental Fig. 12.** Heatmap showing the average total number of IBD segments shared between 3rd level communities in Ireland and the UK and regional European clusters across length bins scaled (z-scores) by column to capture variation in sharing profiles with European clusters across all Irish and British genetic communities.

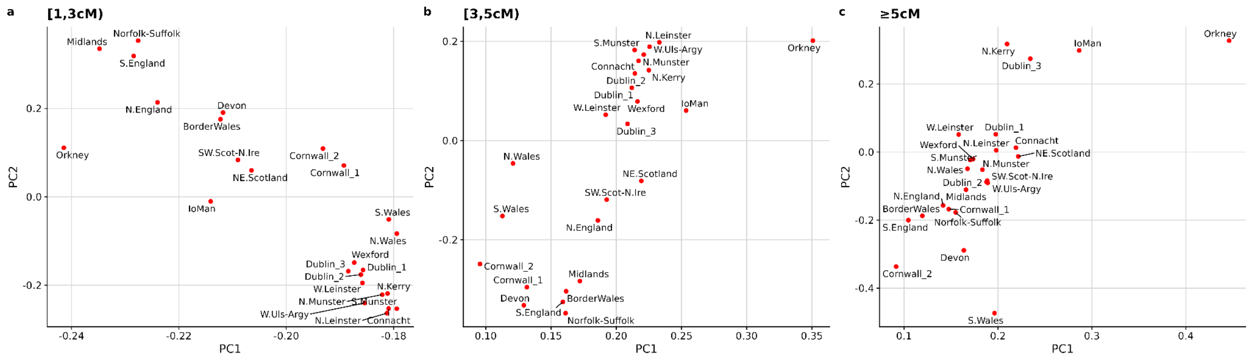


**Supplemental Fig. 13.** Principal components analysis on the average total length of IBD shared between 3rd level communities in Ireland and the UK and regional European clusters generated using fineSTRUCTURE across length bins. These figures show the top 2 principal components.
